# Ancient genomic regulatory blocks are a major source for gene deserts in vertebrates after whole genome duplications

**DOI:** 10.1101/776369

**Authors:** María Touceda-Suárez, Elizabeth M. Kita, Rafael D. Acemel, Panos N. Firbas, Marta S. Magri, Silvia Naranjo, Juan J. Tena, Jose Luis Gómez-Skarmeta, Ignacio Maeso, Manuel Irimia

## Abstract

We investigated how the two rounds of whole genome duplication that occurred at the base of the vertebrate lineage have impacted ancient microsyntenic associations involving developmental regulators (known as genomic regulatory blocks, GRBs). We showed that the majority of GRBs present in the last common ancestor of chordates have been maintained as a single copy in humans. We found evidence that dismantling of the additional GRB copies occurred early in vertebrate evolution often through the differential retention of the regulatory gene but loss of the bystander gene’s exonic sequences. Despite the large evolutionary scale, the presence of duplicated highly conserved non-coding regions provided unambiguous proof for this scenario for dozens of ancient GRBs. Remarkably, the dismantling of ancient GRB duplicates has contributed to the creation of large gene deserts associated with regulatory genes in vertebrates, providing a widespread mechanism for the origin of these enigmatic genomic traits.

## Introduction

Complex spatiotemporal regulation of transcription is crucial for animal embryonic development. This regulation relies heavily on long-range distal enhancers, which are a hallmark of metazoans (Irimia et al. 2013; Sebe-Pedros et al. 2016) and are particularly prevalent in vertebrates (Marlétaz et al. 2018). Long-range enhancers engage in precise physical interactions with their target promoters, which can be located hundreds of kbps away. These enhancers are particularly prevalent in the case of developmental transcription factors (hereafter *trans-dev* genes (Woolfe et al. 2005)), and create complex *cis*-regulatory landscapes that leave distinctive signatures on how the genome is organized around these genes. Among these signatures, the massive size of the intergenic regions associated with *trans-dev* genes is probably the most conspicuous (Nelson et al. 2004). In the most extreme cases, these gene-free regions can be longer than one Mbp, and are commonly known as gene deserts (Nobrega et al. 2003). Most gene deserts associated with *trans-dev* genes are conserved across vertebrates (Ovcharenko et al. 2005), but how they have originated remains a mystery.

In addition to intergenic regions, distal enhancers from *trans-dev* genes can also constrain genomic organization when they are located within the introns of neighboring genes. This creates microsyntenic associations between *trans-dev* and non-*trans-dev* (“bystander”) genes, known as genomic regulatory blocks (GRBs; Figure 1A) (Kikuta et al. 2007). GRBs are thus the expanded regulatory landscapes of the *trans-dev* genes, and closely coincide with topological associated domains (TADs) (Harmston et al. 2017). Importantly, since the disruption of GRBs would separate enhancers from their target *trans-dev* genes, these microsyntenic associations are often highly conserved in evolution (Engstrom et al. 2007; Kikuta et al. 2007; Dong et al. 2009; Irimia et al. 2012b). However, the genetic redundancy created by the whole genome duplication (WGD) of teleosts was shown to have triggered a wholesale dismantling of GRB duplicates (Kikuta et al. 2007; Dong et al. 2009). In this lineage, the most common outcome of the dismantling was the preservation of the *trans-dev* gene and associated enhancers accompanied by the loss of the bystander copy’s exonic sequences (Figure 1A, scenario [ii]).

**Figure 1.**
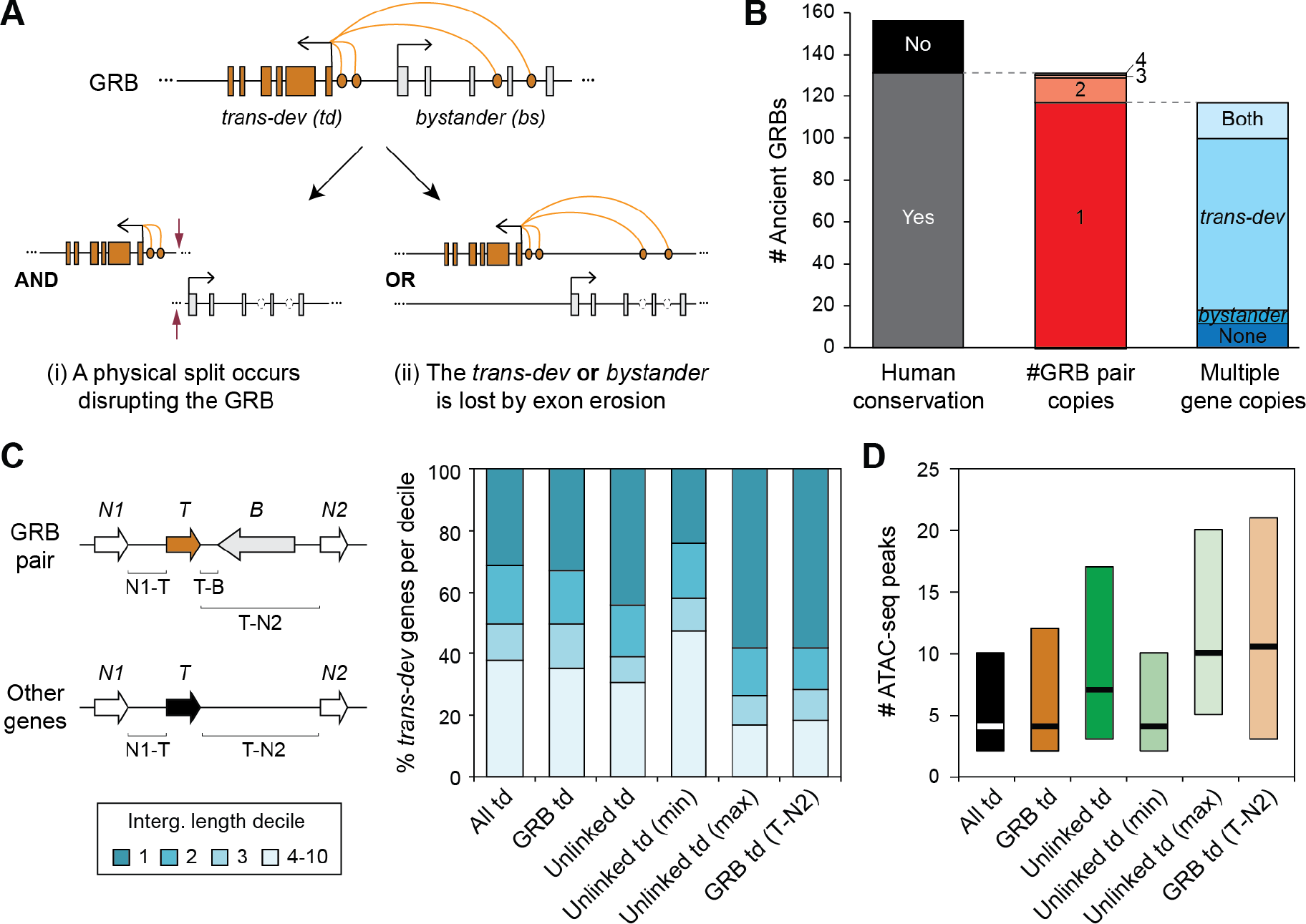
Fates of ancient GRBs after the two WGDs in vertebrates. A) Genetic redundancy allows the dismantling of GRBs after WGD. Two scenarios are depicted: (i) The GRB microsyntenic association is disrupted by a break point, in which both the *trans-dev* and the bystander gene may be maintained, but in different genomic locations; This scenario would impact the regulation of the *trans-dev* gene. (ii) The GRB microsyntenic association is dismantled by the differential loss of either the *trans-dev* or the bystander gene. The loss can occur by a large deletion of the genomic locus or by pseudogenization and “exon erosion”. While the former would impact the regulation of the *trans-dev* when the bystander is lost, the latter would not. B) Summary of the fates of the 156 studied ancient GRB pairs in human. “Human conservation”: whether the human genome has conserved at least one linked copy or not. “#GRB pair copies”: for those conserved, the number of copies of GRB pairs maintained in human (1-4). “ Multiple gene copies”: for those GRB pairs in single copy, in how many cases there are multiple ohnologs for both the *trans-dev* and bystander genes (i.e. not linked), only for the *trans-dev* or bystander gene, or for none. C) Percent of *trans-dev* genes of different types whose intergenic regions are within the first, second, third or another decile genome-wide (i.e. *trans-dev* genes in the first decile have an intergenic region among the top 10% of all genes). “All td”: all *trans-dev* genes linked to at least one non-*trans-dev* gene; “GRB td”: *trans-dev* genes that are part of conserved ancient GRB pairs; “Unlinked td”: *trans-dev* ohnologs that are not linked to the ancient bystander gene. “Unlinked td (max/min)”: the unlinked *trans-dev* ohnolog with the largest/smallest intergenic region. “GRB td (T-N2)”: distance from the *trans-dev* gene in a conserved ancient GRB pair to the gene after the bystander (gene N2). For all categories but “GRB td (T-N2)”, only the largest intergenic region of the two potential ones (N1-T or T-B for linked *trans-dev* genes, or N1-T or T-N2 for the rest) is considered for fairness. D) Number of ATAC-seq peaks found in the intergenic regions for each type of *trans-dev* gene. As in (C), only the intergenic region with the highest number of peaks is considered, except for “GRB td (T-N2)”.

In the case of the two rounds of WGD that occurred at the base of vertebrates (Dehal and Boore 2005; Putnam et al. 2008), it is unknown how they have impacted the evolution of ancient GRBs. Furthermore, except in the case of a few isolated loci (Irimia et al. 2012a; Maeso et al. 2012; Acemel et al. 2016), the mechanisms leading to ancient GRB dismantling are poorly understood. Here, we identified GRBs that were present in the last common ancestor of chordates and investigated their fates after the vertebrate WGDs. We found that the microsyntenic associations of most ancient GRBs have been maintained in single copy, and that the majority of copies dismantled these associations very early in vertebrate evolution, likely by the differential loss of the bystanders and retention of the *trans-dev* genes. Interestingly, we observed that such loss of bystanders have significantly contributed to create multiple human gene deserts, and propose that this is a major source of regulatory gene desert formation at the origin of vertebrates.

## Results

### Evolutionary history of ancient GRBs in the human genome

To investigate the fate of ancient GRBs after the vertebrate WGDs, we assembled a list of putative GRBs that were likely present in the last common ancestor of chordates. We took previously reported microsyntenic non-paralogous gene pairs detected using 13 metazoan species (Irimia et al. 2012b), and also performed a *de novo* search for gene pairs with conserved microsynteny employing more recently published genomes from slow-evolving non-vertebrate species (see Methods for details). From the union of these ancient microsyntenic associations, we then identified those gene pairs containing a developmental transcription factor (*trans-dev*) gene and a non-developmental (bystander) gene. This resulted in a list of 116 ancient putative GRBs. The majority of these GRBs (85/116, 73.3%) contained only one microsyntenic association between a *trans-dev* and a bystander gene, but in the remaining 31 cases the *trans-dev* gene was linked to two or more bystander genes (Supplementary Table S1). Thus, these 116 ancient GRBs corresponded to a total of 156 unique *trans-dev-*bystander ancient syntenic pairs, hereafter GRB pairs. For simplicity, we here studied the evolution of these GRB pairs independently. For specific comparisons, we also compiled a list of 52 ancient human co-regulated non-developmental gene pairs in head-to-head orientation (likely sharing a bidirectional promoter) (Irimia et al. 2012b) (Supplementary Table S2).

The human genome has retained at least one copy of 131/156 (84.0%) ancient GRB pairs (Figure 1B and Supplementary Table S1). However, the microsyntenic association between *trans-dev* and bystander was maintained in more than one copy for only 14/131 (10.7%) of those ancestral GRB pairs (e.g., ONECUT1/WDR72 on chromosome 15 and ONECUT2/WDR7 on chromosome 18) (Figure 1B and Supplementary Table S1). A similar percentage of co-regulated gene pairs (3/52, 5.8%) were also maintained in multiple copies (p = 0.41, two-sided Fisher’s exact test)(Supplementary Table S2). The large fraction of single-copy gene associations after WGD could be due to several reasons. First, even if multiple ohnologs (i.e. paralogs derived from WGDs) are maintained for both the *trans-dev* and the bystander genes, their microsyntenic association may be lost by genomic rearrangements (Figure 1A, scenario [i]). Second, *trans-dev* and non-*trans-dev* genes are known to be retained at different rates after WGD (Putnam et al. 2008; Cañestro et al. 2013), which could also result in the loss of the GRB syntenic association (Figure 1A, scenario [ii]). Consistent with the second scenario, only 17/117 (14.5%) of single-copy GRB pairs have retained multiple copies of both the *trans-dev* and the bystander gene (Figure 1B and Supplementary Table S1). In the vast majority of cases, only the *trans-dev* gene has been retained in multiple copies (82/117, 70.1%), compared to 7 cases (6.0%) for the bystander gene (p = 8.57e-12, proportion test). These differences are due to higher retention rates of the *trans-dev* genes, since the bystander genes were retained in multiple copies at rates similar to those of genes in co-regulated pairs (24/117 [20.5%] vs. 26/104 [25%]; P = 0.54, two-sided Fisher’s exact test). The differential loss of bystander genes is illustrated by the cases of ancestral multi-bystander GRBs in which different *trans-dev* ohnologs have remained associated with only one of the bystander families (Supplementary Figures S1 and S2 and Supplementary Table S1). In these cases, it is likely that specific bystanders were differentially lost in each vertebrate GRB copy, similar to what has been reported in teleosts (Kikuta et al. 2007; Dong et al. 2009).

To determine the relative timing at which the loss of the GRB syntenic associations occurred, we next studied ancestral GRB pair evolution in two slow-evolving basal-branching vertebrate species, the elephant shark *Callorhinchus milii* and the spotted gar *Lepisosteus oculatus*, and in chicken. We could find evidence of linkage for additional pairs of *trans-dev* and bystander ohnologs that are not linked in the human genome for only 2/131 (1.5%) ancestral GRB pairs: *FOXP3/GPR173* in *L. oculatus*, and *PBX4/MVB12A* in *L. oculatus*, *C. milii* as well as in chicken (Supplementary Table S1). A similar pattern was found for the 25 ancestral GRB pairs that have not been conserved in human, for which only two (8%) gene pairs were linked in any of the other vertebrate genomes: *MKL2/DCUN1D3* in *C. milii*, and *KDM1A/TMEM30B* in *C. milii* and *L. oculatus*. Therefore, these data indicate that the vast majority of losses of GRB syntenic associations occurred before the last common ancestor of gnathostomes, soon after the two rounds of WGD.

### Dismantlement of ancient GRB pairs often contributes to gene deserts in the human genome

Our results thus far show that the majority of ancient GRB pair associations have been conserved in a single copy and that most losses occurred early in vertebrate evolution, likely involving the differential loss of the bystander copies and preservation of *trans-dev* ohnologs (scenario [ii] in Figure 1A). The differential loss of the bystander gene could occur by a large deletion of the locus or by pseudogenization of the exonic sequence (“exon erosion”), with dramatically different effects on the conservation of the regulatory landscape of the *transdev* gene: whereas a large deletion would remove putative regulatory elements of the *transdev* gene, erosion of the *bystander* exons would allow essential regulatory elements to be maintained. Interestingly, considering that bystanders are usually very large genes (Kikuta et al. 2007; Irimia et al. 2012b), the latter could result in the creation of a large gene-free region in the formerly-bystander locus, i.e. a “gene desertification” of the GRB.

To evaluate the predictions made by this hypothesis, we first calculated the length of the intergenic regions around different sets of *trans-dev* genes in the human genome (Supplementary Table S3). As expected (Nelson et al. 2004), *trans-dev* genes that are part of GRBs with conserved ancient synteny (“GRB td”) were enriched for large intergenic regions, but to an extent similar to that of the whole set of human *trans-dev* genes associated with at least one non-*trans-dev* gene (Figure 1C; 33.0% of “GRB td” are in the top decile of intergenic distances genome-wide (>229 Kbp), and 65.1% among the top three deciles, compared to 31.2% and 62.2% for all *trans-dev* genes [“All td”]; p=0.7361, Fisher’s Exact test for the first deciles). The fraction of top deciles was significantly increased for the case of the *trans-dev* ohnologs from ancient GRB pairs that are no longer linked to the original bystander genes (“Unlinked td”; 44.4% and 69.6%, p = 0.0013, Fisher’s Exact test compared to all *trans-dev* genes). Interestingly, this enrichment was much stronger when only the unlinked *trans-dev* ohnolog with the largest intergenic region was considered (“Unlinked td (max)”, 58.3% and 81.6%, p = 1.36e-07, Fisher’s Exact test), consistent with the asymmetric evolution of gene regulatory landscapes reported for vertebrate ohnologs (Marlétaz et al. 2018). Remarkably, this distribution of intergenic distances closely matches that of the distances between the *trans-dev* genes in conserved ancestral GRB pairs and the next neighboring gene after the bystander (i.e. including the bystander, “GRB td (T-N2)”; Figure 1C). Similar patterns were observed when comparing the complexity of regulatory landscapes among sets of *trans-dev* genes, measured as the number of ATAC-seq peaks found in the different intergenic regions (Figure 1D). For this, we used ATAC-seq data for multiple stages from twelve developing mouse tissues from the ENCODE project, and counted the number of significant ATAC-seq peaks detected in the mouse orthologous regions in at least two tissues (see Methods for details). Consistent with the differences in intergenic region lengths, unlinked *trans-dev* genes from conserved ancestral GRB pairs have higher numbers of associated ATAC-seq peaks compared to linked (and all) *trans-dev* genes (Figure 1D). This is even higher for the unlinked ohnolog with the largest number of peaks (“Unlinked td (max)”), whose distribution is again similar to that observed for linked *trans-dev* genes when the bystander loci are considered as part of the *trans-dev* genes’ intergenic region (“GRB td (T-N2)”).

These patterns are thus consistent with at least a significant fraction of ancestral GRB pairs having been dismantled by exon erosion, keeping a similarly large and complex regulatory landscape associated with the *trans-dev* gene. However, the erosion of *bystander* exons in duplicated GRBs can only be unequivocally demonstrated by detecting *bystander* pseudogenic exon remnants or by the presence of duplicated highly conserved non-coding regions (HCNRs) that were originally located within the introns of the *bystander* gene before the two WGDs (McEwen et al. 2006; Wang et al. 2007; Maeso et al. 2012; Acemel et al. 2016). To investigate these possibilities, we used a set of duplicated HCNRs in which one of the copies is present within a bystander gene of an ancient GRB pair and the other in a gene-free region surrounding the paralogous *trans-dev* gene, which we compiled from a previous report (McEwen et al. 2006) and from a *de novo* search using mouse embryonic ATAC-seq peaks (see Methods). In total, we identified a total of 23 pairs of HCNRs associated with 16/99 (16.2%) ancestral GRB pairs in which at least one linked and one unlinked *trans-dev* genes have been retained in human (Supplementary Table S4). In addition, we performed a systematic search for exon remnants from bystanders in the intergenic regions of unlinked *trans-dev* genes, which resulted in the discovery of a high-confidence exon remnant from a paralog of *FAM172A* associated with *NR2F2* (Supplementary Table S4).

One example of bystander erosion is provided by the GRB pair formed by *Islet/Scaper*, conserved from human to sponges (Irimia et al. 2012b; Wong et al. 2019). McEwen *et al*. identified two duplicated HCNRs in mammals, one in the intron of *Scaper*, near *Isl2*, and one in a gene desert of 1.4 Mbp near *Isl1* (McEwen et al. 2006). In both cases, *Isl1* and *Isl2*, along with the conserved duplicated elements, were located within the borders of single mouse TADs (Figure 2A). These two sequences were conserved across multiple vertebrates, including zebrafish (Figure 2B). To assess if the duplicated HCNRs interacted with the respective Islet promoters, we generated circular chromosome conformation capture sequencing (4C-seq) using zebrafish *isl1* and *isl2a* promoters as viewpoints. In both cases, the HCRNs interacted with the respective *Islet* promoters (Figure 2B). Furthermore, we generated 4C-seq data in amphioxus embryos using the promoter of its single *Isl* gene as viewpoint, and found it to interact with the orthologous Scaper locus (Figure 2B). Next, to probe the potential enhancer activity of these elements, we generated stable zebrafish lines using the ZED vector (Bessa et al. 2009). Both HCNRs drove expression in the nervous system in overlapping, but also distinct domains, consistent with the expression of their target genes (Figure 2B,C). A similar scenario was observed for other *trans-dev* genes with large gene deserts that were originally involved in ancestral GRBs, such as *Otx1/2-Ehbp1*, for which we could also show interaction between duplicated HCNRs and their respective promoters in zebrafish and between *Otx* and the orthologous *Ehbp1* locus in amphioxus (Supplementary Figure S3).

**Figure 2.**
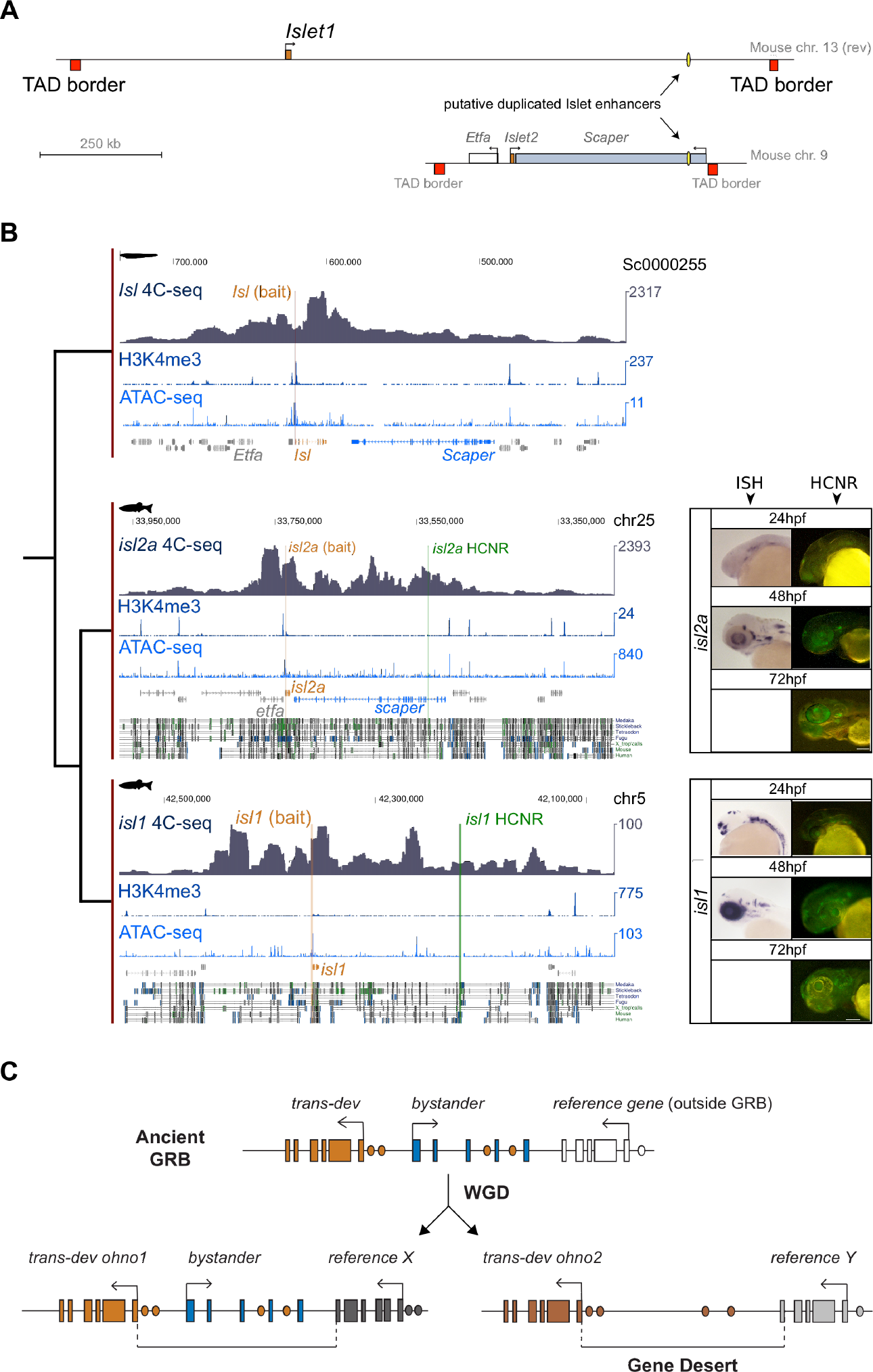
The evolution of the *Islet-Scaper* GRB pair exemplifies the contribution of bystander erosion to the origin of gene deserts. A) Schematic representation of *Isl1* and *Isl2* gene regulatory landscapes in the mouse genome (regions contained between the TAD borders identified by (Dixon et al. 2012)). *Scaper*, the bystander associated with *Isl2*, is depicted, as well as the pair of ohnologous HCNRs identified by (McEwen et al. 2006). **B)** 4C-seq signal using *Isl* promoters as viewpoints (orange) and H3K4me3 and ATAC-seq signal from 24 hpf zebrafish or 15 hpf amphioxus embryos, and conservation tracks from the UCSC Genome Browser (for zebrafish only); ohnologous HCNRs are highlighted in green. Right: *In situ* hybridization of *isl1* and *isl2a* from (Thisse et al. 2004) and GFP expression driven by the HCNRs associated with *isl1* and *isl2a* at different timepoints using the ZED vector. During the first few days of development, the *isl1* enhancer showed dynamic expression. At 24 hpf the reporter shows expression in the anterior telencephalon, and a subset of retina and ventral hindbrain cells. At 48 and 72 hours, it maintains expression in a sparse subset of neural cells that may overlap with small, localized regions of *isl1* expression. In the case of *isl2a*, the enhancer found within *scaper* drove consistent expression in a diffuse anterior domain at 24 hpf, as well as in the pineal gland and ventral hindbrain. The expression becomes more restricted at 48 hpf and 72 hpf; to the pineal gland, subsets of the retinal and otic cells, and faintly in the diencephalon. Scale bar: 100 μm. C) Model of gene desert formation by GRB dismantling through bystander exon erosion.

## Discussion

We found that most ancient GRB pairs have been retained in a single copy in the vertebrate lineage after two rounds of WGD. Only a small subset of GRB pairs have been maintained in multiple copies, a fraction similar to that observed for ancient pairs of co-regulated genes in a head-to-head orientation, suggesting no bias for or against retention of GRB syntenic associations in multiple copies after WGD. Moreover, data from slow-evolving, basal-branching gnathostomes showed that the dismantlement of extra duplicates likely occurred shortly after the two WGDs. Although it is difficult to assess it confidently given the amount of evolutionary time involved, we also found evidence that a significant fraction of the dismantled GRB syntenic associations happened by differential loss of the bystander gene by pseudogenization (“bystander exon erosion”). This pattern is similar to that reported for the more recent teleost-specific WGD (Kikuta et al. 2007; Dong et al. 2009), for which a larger number of HCNRs within bystander genes could be used to unequivocally prove this fate. In the case of the vertebrate WGDs, a few pre-duplicative HCNRs provided such proof for around 16% of the ancient GRB pairs. Given that sequence conservation of ancient HCNRs is extremely rare (McEwen et al. 2006; Royo et al. 2011), it is possible that similar processes have occurred with many other ancient GRBs in vertebrates despite the lack of conservation of duplicated HCNRs, as suggested by the analysis of intergenic length distributions. Remarkably, for many cases in which a *trans-dev* ohnolog has been retained without the association with a bystander copy, we could observe an unusually large associated gene-free region, including for those ones with HCNR-based evidence for bystander erosion. This highlights a mechanism for gene desert creation after WGD (Figure 2D), by which the already-large intergenic regions of *trans-dev* genes are massively increased by the addition of the genomic sequences corresponding to the bystander loci. This is in contrast to a gradual increase of gene-free regions through evolutionary time as the major path for gene desert formation, although both scenarios may certainly occur together. Importantly, although this model is based on the data from GRB pairs present in the last common ancestor of chordates, and may thus explain only a small fraction of human gene deserts, it should be noted that we do not know the actual repertoire of GRB syntenic associations present immediately before the vertebrate WGDs. Therefore, many or even most vertebrate gene deserts associated with *trans-dev* genes (known as regulatory gene deserts (Ovcharenko et al. 2005)) might have originated by desertification of pre-duplicative GRBs. Whatever the extent, disintegration of bystanders from duplicated GRBs certainly contributed to explain the origin of this enigmatic genomic trait, which is particularly prevalent in vertebrates.

## Materials and Methods

We assembled a comprehensive catalog of ancient GRBs (present in the last common ancestor of chordates) using three main sources: (i) gene pairs with ancient microsyntenic associations identified by comparing 13 metazoan genomes (Irimia et al. 2012b) (595 pairs); (ii) gene pairs containing duplicated HCNRs identified by (McEwen et al. 2006) (18 pairs); and (iii) a *de novo* search for ancient microsyntenic associations (1,538 pairs; Supplementary Table S5, see Supplementary Materials and Methods for details). Gene pairs from the three sources were then combined into a non-redundant set of pairs, and we defined ancient GRB pairs as gene pairs formed by a *trans-dev* and a non-*trans-dev* gene. *trans-dev* genes were defined as “transcription factors involved in the regulation of developmental processes” based on Gene Ontology information.

To investigate the evolution of GRB pairs after WGDs, we first defined a consensus set of ohnologs based on different sources ((Makino and McLysaght 2010; Singh et al. 2015) and Ensembl Paralogs)(Supplementary Table S6). Based on this ohnology information, for each tested ancestral GRB, we counted the number of *trans-dev* and bystander ohnologs conserved in the human genome and re-assessed which pairwise combinations were linked together and separated by no more than two intervening genes. Genes that had no ohnolog were assumed to have gone back to single copy after the two WGDs.

To obtain pairs of ohnologous regulatory elements that were differentially associated with ohnologous *trans-dev* genes (one in a bystander and another one in an intergenic region), we collected putative ancient duplicated enhancers from two sources: (i) duplicated HCNRs identified by (McEwen et al. 2006) and reported to be within a neighboring gene and an intergenic region; and (ii) a *de novo* search for duplicated ATAC-seq-defined regulatory elements. For the latter, we downloaded 132 ATAC-seq experiments corresponding to several stages from twelve mouse developing tissues (biosamples) from the ENCODE portal (Supplementary Table S7)(see Supplementary Materials and Methods for details).

The length of the intergenic region between each gene pair was calculated for the human hg38 genome, using only one representative transcript per gene, and considering only protein-coding genes that were included in the OrthoFinder homology clusters. Telomere and centromere regions (downloaded from the UCSC Table function) were discarded. Intergenic regions were then ranked and deciles were made (the top decile corresponded to regions of at least 228,558 bps, and the first three deciles to 54,957 bps; the median intergenic region was 19,949 bps).

4C-seq experiments were performed and analyzed as described earlier (Acemel et al. 2016)(primers provided in Supplementary Table S8). To probe the conservation of HCNRs associated with *isl2a* and *isl1* in zebrafish, and the open chromatin region located between *hey2* and *hddc2* (*hey2-E1)*. The sequences were cloned into the Zebrafish Enhancer Detection (ZED) vector and transgenic lines were generated and screened as previously described (Bessa et al. 2009).

## Acknowledgements

The research has been funded by the European Research Council (ERC) under the European Union’s Horizon 2020 research and innovation program (ERC-StG-LS2-637591 to M.I. and ERC-AdG-LS8-740041 to J.L.G.-S), the Spanish Ministerio de Economía y Competitividad (BFU2017-89201-P to M.I., RYC-2016-20089 to I.M., BFU2016-74961-P to J.L.G.-S., and BFU2016-81887-REDT/AEI to J.L.G.-S and M.I.), the People Programme (Marie Curie Actions) of the European Union's Seventh Framework Programme (FP7/2007-2013) under REA grant agreement 608959, the ‘Centro de Excelencia Severo Ochoa 2013-2017’(SEV-2012-0208) and the ‘Unidad de Excelencia María de Maetzu 2017-2021’(MDM-2016-0687). We acknowledge the support of the CERCA Programme/Generalitat de Catalunya and of the Spanish Ministry of Economy, Industry and Competitiveness (MEIC) to the EMBL partnership.

## Author Contributions

M.T.-S., R.D.A., P.N.F., I.M. and M.I. performed bioinformatic analyses. E.M.K., R.D.A., M.M., S.N., J.J.T performed molecular biology and transgenesis experiments. J.L.G.-S., I.M. and M.I. conceived, designed and coordinated the study. E.M.K., I.M. and M.I. wrote the manuscript with the input of the other authors.

