## Supplementary Data for "Ancient genomic regulatory blocks are a major source for gene deserts in vertebrates after whole genome duplications"

### Supplementary Figures

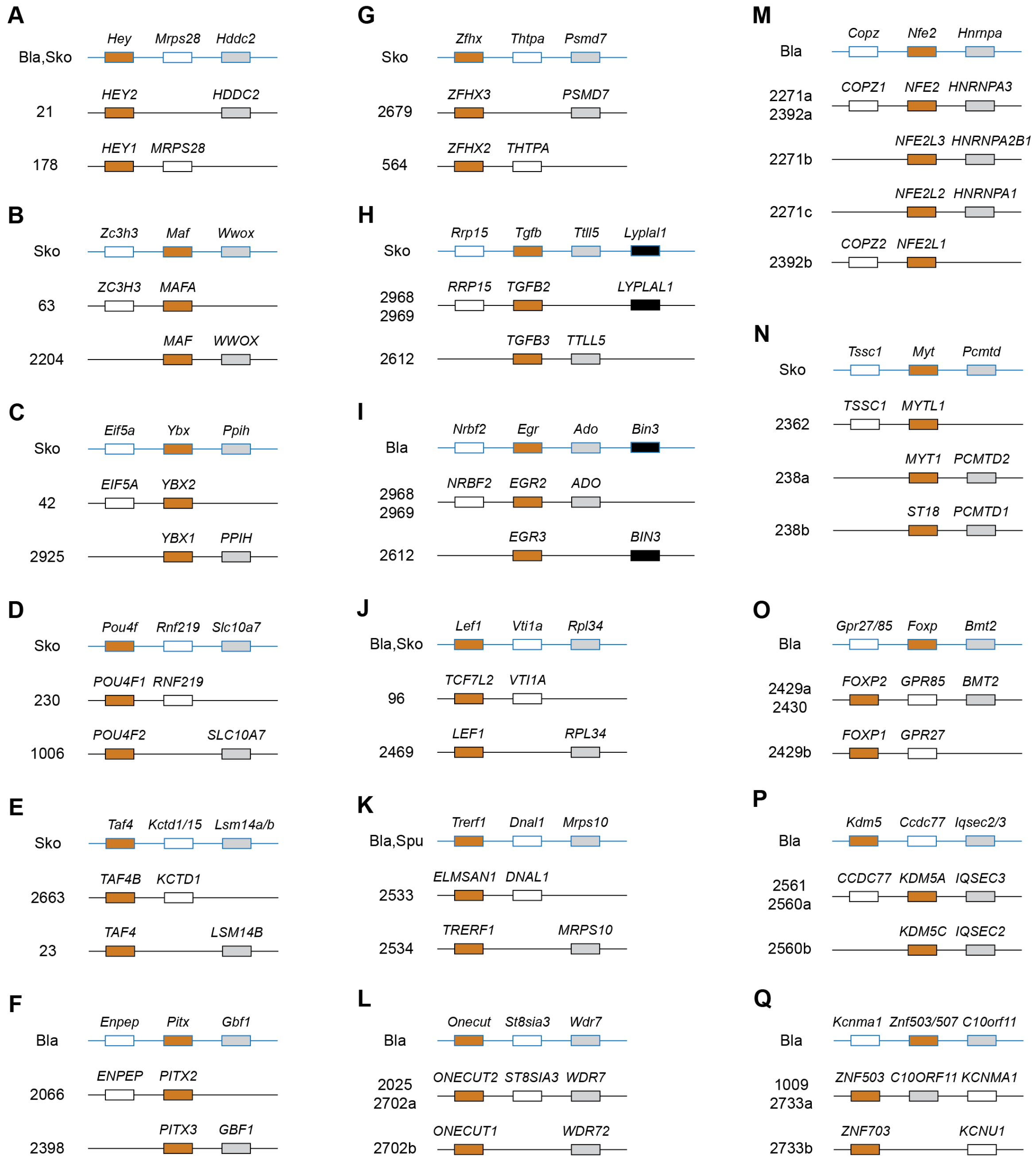


**Supplementary Figure S1 - Microsyntenic arrangements of ancient multi-bystander GRBs whose bystanders have become differentially retained next to different *trans-dev* ohnologs.** For each case, the arrangement in a slow-evolving deuterostome (Bla, *B. lanceolatum*; Sko, *S. kowalevskii;* Spu, *S. purpuratus*) is provided on top (blue lines), followed by the GRB arrangements conserved in the human genome.

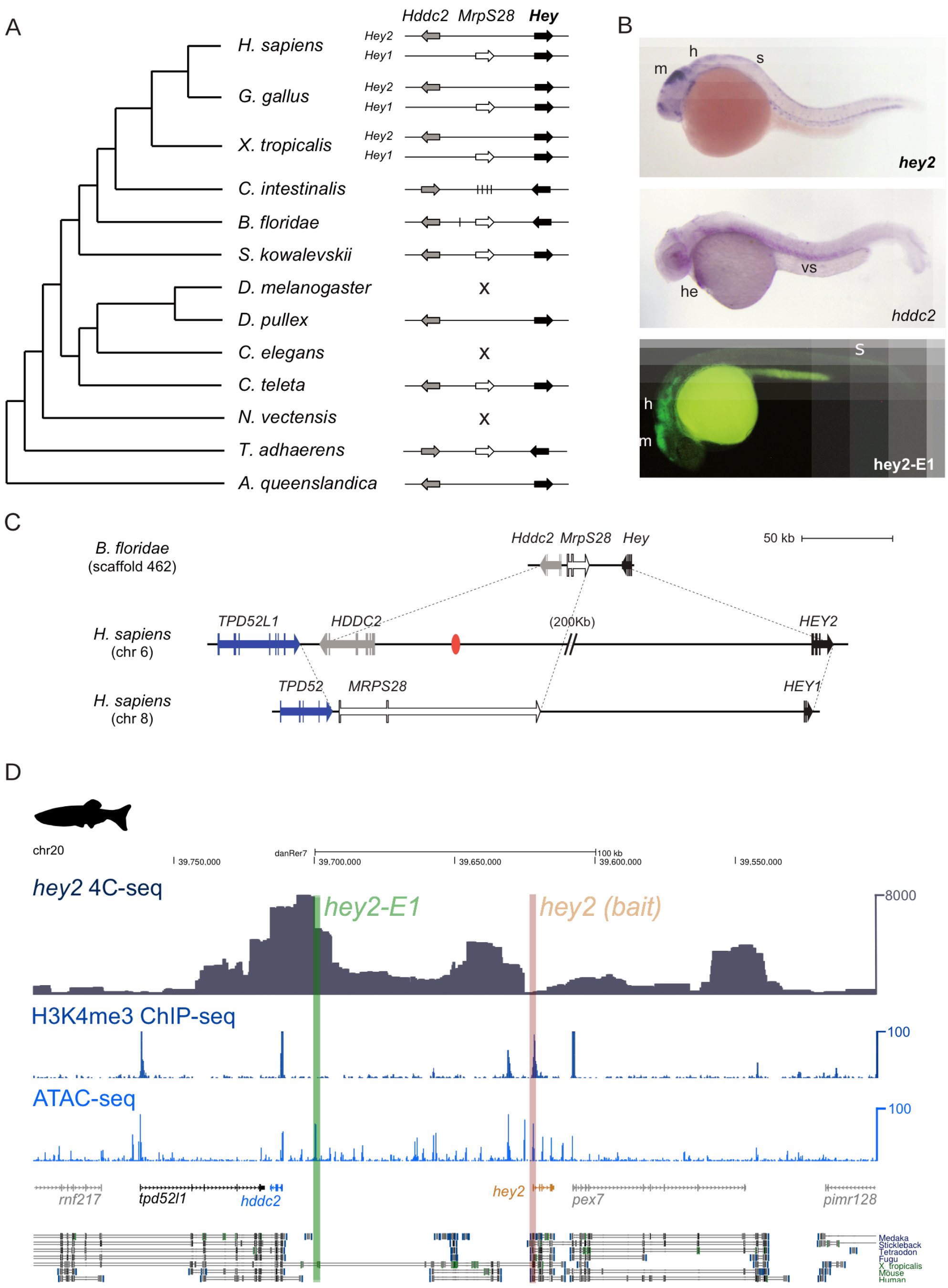


**Supplementary Figure S2 - Evolution of the *Hey-MrpS28-Hdcc2* GRB and its functional characterization in zebrafish.** A) Phylogenetic distribution of the GRB across the studied metazoan species. Only *B. floridae*, *S. kowalevskii*, *C. teleta* and *T. adhaerens* have conserved both *bystander* genes: *Hddc2* (grey) and *MrpS28* (white), linked to the *trans-dev* gene *Hey* (black arrows). In vertebrates, each of the Hey paralogs has preserved only one of the *bystander* genes in a reciprocal manner. No data were available for *S. purpuratus* or *L. gigantea*. B) Upper and middle panels are zebrafish embryos at 24 hpf showing the expression of *hey2* and *hddc2* genes. *hey2* is detected in the dorsal midbrain (m), hindbrain (h) and spinal cord (s) while *hddc2* is detected in the heart (he). Lower panel shows GFP expression promoted by an enhancer located close to the *hddc2* gene (green rectangle in D). This enhancer is active in embryonic domains expressing *hey2*. C) Syntenic organization at scale and gene structures of the GRBs in the cephalochordate *B. floridae*, with both *Hddc2* (grey) and *MrpS28* (white)

bystanders, and both human *HEY* paralogs. The ohnologous pair *TDP52* and *TDP52L1* delimits both vertebrate GRB copies, supporting that the differential losses of bystanders was due to gene loss and not to chromosomal rearrangements. The red ellipse indicates the approximate position (~44 kbp to *HDDC2* and ~404 kbp to *HEY2*) of the orthologous region to the sequence with positive enhancer activity and physical interaction with the *hey2* promoter in zebrafish (green rectangle in D). D) Distribution of H3K4me3 and ATAC-seq signal in the *hey2-hddc2* GRB, and 4C-seq signal using the *hey2* promoter as viewpoint in 24 hpf zebrafish embryos. An green rectangle indicates the tested regulatory element, which shows a strong contact with *hey2*.

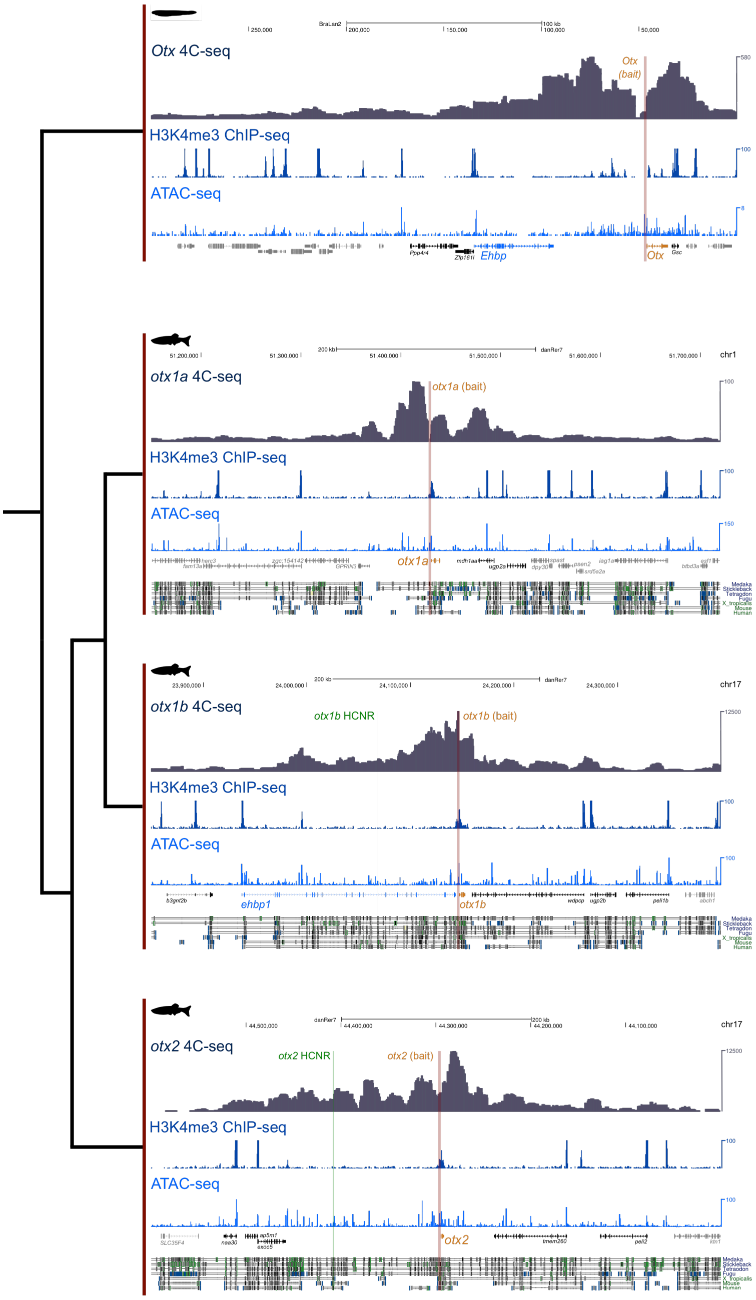


**Supplementary Figure S3 - Evolution of the *Otx-Ehbp1* GRB pair.** 4C-seq signal using *Otx* promoters as viewpoints (orange) and H3K4me3 and ATAC-seq signal from 24 hpf zebrafish or 15 hpf amphioxus embryos, and conservation tracks from the UCSC Genome Browser (for zebrafish only). The two ohnologous HCNRs identified by (McEwen, et al. 2006) are highlighted in green.

**Supplementary Methods**

***Identification and definition of ancient GRBs***

We assembled a comprehensive catalog of ancient GRBs (present in the last common ancestor of chordates) using three main sources: (i) gene pairs with ancient microsyntenic associations identified by comparing 13 metazoan genomes (Irimia, et al. 2012) (595 pairs); (ii) gene pairs containing duplicated HCNRs identified by (McEwen, et al. 2006) (18 pairs); and (iii) a *de novo* search for ancient microsyntenic associations (1,538 pairs). For the latter, we employed the following genome assemblies and annotations: human (*Homo sapiens*, hg38 Ensembl v88), chicken (*Gallus gallus*, galGal4 Ensembl v83), spotted gar (*Lepisosteus oculatus*, LepOcu1 Ensembl v88), elephant shark (*Callorhinchus milii*, ESHARK1 (Venkatesh, et al. 2014)), amphioxus (*Branchiostoma lanceolatum*, Bl71nemr (Marlétaz, et al. 2018)), acorn worm (*Saccoglossus kowalevskii*, Skow_1.0, https://groups.oist.jp/molgenu/hemichordate-genomes), sea urchin (*Strongylocentrotus purpuratus*, Spur3.1, http://www.echinobase.org/Echinobase/), centipede (*Strigamia maritima*, Smar1 Ensembl Metazoans v26) and limpet (*Lottia gigantea*, Lotgi1 Ensembl Metazoans v26). We next selected a single representative transcript per gene for each species, and ran OrthoFinder v2.2.7 (Emms and Kelly 2015) to produce gene homology relationships among the nine species with the following parameters: -M msa -T raxml-ng -I 1.3 -S diamond.

We then performed pairwise comparisons to identify conserved microsyntenic associations between pairs of species. In brief, for any pair of neighboring genes in species 1, we required orthologs of both genes to be separated by no more than one intervening gene in species 2. Pairs of paralogs genes (i.e. belonging to the same OrthoFinder group) were discarded. Also, if either of the two genes in a pair belonged to an OrthoFinder group with more than 10 members (considering only the two species under comparison), the pair was discarded. This process was repeated for each pair of species using the pairwise OrthoFinder output, as well as the merged OrthoFinder groups for the nine species. The number of pairs identified for each pair of species is provided in Supplementary Table S5. Next, we combined this information to identify pairs of genes that were likely linked since the last common ancestor of chordates. For this, we defined three possibilities: pairs associated in at least (i) two vertebrate species and amphioxus (724 pairs), (ii) two vertebrates and two non-chordates (301 pairs), or (iii) amphioxus and two non-chordates (513 pairs).

The gene pairs from the three sources were then combined into a non-redundant set of pairs, and we defined ancient GRB pairs as gene pairs formed by a *trans-dev* and a non-*trans-dev* gene. *trans-dev* genes were defined as "transcription factors involved in the regulation of developmental processes". For this purpose, we queried human GO terms from Biomart Ensembl v94 as follows:

- Development: contains the GO term GO:0030154 (cell differentiation) or at least two GO terms whose names contain the word "development" (e.g. brain development or embryo development).
- Transcription factor: contains the GO term GO:0043565 (sequence-specific DNA binding), or at least one GO term whose name contains the word "transcription" and contains at least one GO term whose name contains "DNA binding".

Moreover, if a given gene is considered a *trans-dev* gene, all its ohnologs (see below) are defined as *trans-dev* as well. For practical reasons, to study the evolutionary fate of GRBs in vertebrates, we considered each *trans-dev*-bystander syntenic pair individually, even if in some cases these syntenic associations are part of a multi-bystander GRB associated with a single *trans-dev* gene. Finally, for a comparison set, we selected 52 co-regulated blocks in a head-to-head (or 5′–5′) orientation with the highest co-expression from (Irimia, et al. 2012).

***Determination of the fate of ancient GRBs and co-regulated pairs in human***

We first compiled a list of all human ohnologous relationships (paralogs resulting from WGDs). For this purpose, we combined three different sources: (i) ohnologs from http://ohnologs.curie.fr/ (Singh, et al. 2015); (ii) ohnologs reported by (Makino and McLysaght 2010); and (iii) human paralogs with "Vertebrata" or "Euteleostomi" ancestry, from Ensembl Biomart v93. To reduce false positives, we considered pairs of ohnologs those reported by at least two sources (using the "Relaxed" set from (Singh, et al. 2015)), or in the "Intermediate" set from (Singh, et al. 2015). Pairs supported by two or fewer sources were manually curated upon inspection of gene symbols. Specific developmental gene super-families (e.g. Pax, Sox, Wnt and Hox) were also manually curated based on literature information. In total, a set of 2,765 ohnolog gene families was defined, including 7,766 genes (Supplementary Table S6).

Based on this ohnology information, for each tested ancestral GRB, we counted the number of *trans-dev* and bystander ohnologs conserved in the human genome and re-assessed which pairwise combinations were linked together and separated by no more than two intervening genes. Genes that had no ohnolog were assumed to have gone back to single copy after the two WGDs.

***Determination of the fate of ancient GRBs in other vertebrates***

Using OrthoFinder orthology information, for each ancestral GRB pair, we first retrieved all assigned orthologs for *trans-dev* and bystander copies that were not linked in the human genome and evaluated their microsyntenic association in the chicken, spotted gar and elephant shark genomes. For those cases in which a *trans-dev* and a bystander copy were detected to be together separated by no more than two intervening genes, we manually inspected the association and orthology assignments, to ensure that no orthology mis-assignments were done and that the copy of the GRB pair conserved in human was also present in these species.

***Identification of ancient duplicated HCNRs and bystander exon remnants***

To obtain pairs of ohnologous regulatory elements that were differentially associated with ohnologous *trans-dev* genes (one in a bystander and another one in an intergenic region), we collected putative ancient duplicated enhancers from two sources: (i) duplicated HCNRs identified by (McEwen, et al. 2006) and reported to be within a neighboring gene and an intergenic region; and (ii) a *de novo* search for duplicated ATAC-seq-defined regulatory elements. For the latter, we downloaded 132 ATAC-seq experiments corresponding to several stages from twelve mouse developing tissues (biosamples) from the ENCODE portal (Supplementary Table S7). Raw ATAC-seq reads were mapped to the mouse genome (mm10 assembly) using bowtie2 (Langmead and Salzberg 2012) with the following options: --very-sensitive -X 2000 -I 0. Read pairs with a fragment length < 120 bp were considered "nucleosome free" and subsequently used for peak calling. For this, MACS2 (Zhang, et al. 2008) was used for low-thresholded peak calling using the following parameters: callpeak --nomodel --keep-dup 1 --llocal 10000 --extsize 74 --shift -37  -p 0.07. Next, the IDR framework (IDR 2.0.3; https://github.com/nboley/idr) was used to obtain high confidence peaks based on replicate information for each stage for each developing tissue. Then, for each tissue, we produced a list of significant ATAC-seq peaks that were identified in at least two stages employing bedtools merge. Finally, a single list of ATAC-seq peaks was produced by selecting those peaks that were detected in at least two developing tissues (biosamples), which was used for further analyses (identification of ohnologous regulatory elements and measurement of gene regulatory complexity of different sets of *trans-dev* genes).

To identify potentially ancient duplicated regulatory elements, we blasted all merged ATAC-seq peaks against each other using the following parameters: blastall -p blastn -e 10 -F F -v 10 -b 10. Next, for each significant hit, we first tested if the query peak fell near (within three genes at each side) any ortholog of the *trans-dev* genes involved in ancient GRBs conserved in humans. If so, we searched for all target ATAC-seq peaks if they also fell nearby any of the *trans-dev* ohnologs. This analysis retrieved 25 potential candidate pairs of ATAC-seq peaks, which were manually investigated. This allowed the identification of three additional pairs of HCNRs (in *Tbx2/3*, *Meis1/2* and *Pax1/9*).

To identify potential eroded bystander exon remnants associated with unlinked *trans-dev* genes, we first extracted the exonic sequences of each linked bystander gene for those ancient GRB pairs with at least one linked and one unlinked *trans-dev* copy. These exonic sequences were blasted against the hg38 genome, using loose conditions (-e 100 -F F). The hits for each exon were first processed to select those in the neighboring region of the corresponding unlinked *trans-dev* gene (up to two neighboring genes at each side). From these, 28 unique hits of length ≥ 25 nucleotides were manually investigated. In addition to reciprocal hits corresponding to paralogous bystanders from GRB pairs retained in two or more copies (e.g. *NFE2/L2/L3*, *CDH7/8*, *KDM2A/FBXL19*, etc.), which served as positive controls of the methodology, only a 50 bp hit corresponding to the 3'UTR of *FAM172A*, the bystander gene of *NR2F1*, passed this filter and was located in the expected relative position downstream of *NR2F2*. This hit was further validated by back-blast. Both the 3'UTR exon of *FAM172A* and its paralogous copy in the *NR2F2* associated desert overlapped with highly conserved non-coding elements across vertebrate species.

***Analysis of intergenic region length and complexity of regulatory landscapes***

The length of the intergenic region between each gene pair was calculated for the human hg38 genome, using only one representative transcript per gene, and considering only protein-coding genes that were included in the OrthoFinder homology clusters. Telomere and centromere regions (downloaded from the UCSC Table function) were discarded. Intergenic regions were then ranked and deciles were made (the top decile corresponded to regions of at least 228,558 bps, and the first three deciles to 54,957 bps; the median intergenic region was 19,949 bps). To define the set of "All td" (Figure 1C,D), all *trans-dev* genes that had at least one non-*trans-dev* gene nearby were selected. For fairness, the largest of the two intergenic regions was used in the plots for each *trans-dev* gene, except for the specified T-B and T-N2 distances for linked *trans-dev* in ancient GRB pairs (see Figure 1C for an schematic representation). To define the complexity of regulatory landscapes, we counted the number of putative regulatory elements, as defined by ATAC-seq, in the corresponding mouse orthologous regions. As before, the region with the largest number of ATAC-seq peak was used for each *trans-dev* gene, unless otherwise specified (i.e. T-B and T-N2).

***4C-seq experiments in amphioxus and zebrafish embryos***

4C-seq experiments were performed and analyzed as described earlier (Acemel, et al. 2016). For the amphioxus samples, around 4000 whole embryos at the stage of 15 h.p.f were collected and fixed for 15 min at room temperature in MOPS buffer (0.1 M MOPS pH 7.5, 2 mM MgSO2, 1 mM EGTA and 0.5 M NaCl) supplemented with PFA with a final concentration of 1.85%. For the zebrafish samples, 500 embryos at the 24hpf stage were dechorianated with pronase and deyolked using the Ginzburg Fish Ringer buffer (55 mM NaCl, 1.8 mM KCl and 1.25 mM NaHCO3). Then they were fixed using 2% PFA for 15 minutes at room temperature. From this step onwards, the 4C-seq protocols are equivalent for both species. Fixations were stopped by adding glycine and washing several times with PBS (NaPBS in the case of amphioxus). Then, the fixed samples were lysed (lysis buffer: 10 mM Tris-HCl pH 8, 10 mM NaCl, 0.3% Igepal CA-630 (Sigma-Aldrich, I8896) and 1× protease inhibitor cocktail (Complete, Roche, 11697498001)), and the DNA was digested with DpnII (New England BioLabs, R0543M) and Csp6I (Fermentas, Thermo Scientific, FD0214) as primary and secondary enzymes, respectively. T4 DNA ligase (Thermo scientific, EL0014) was used for both ligation steps. Specific 4C-seq primers containing single-end Illumina adaptors were designed using as bait the different promoters of interest (see Supplementary Table S8). For each library, eight independent PCRs were performed using the Expand Long Template PCR System (Roche, 11759060001) that were subsequently pooled and purified using AMPure XP beads (Beckman Coulter, A63883). Then libraries were sent for single-end sequencing.

4C-seq sequencing reads were aligned to the reference genomes of either *Danio rerio* (danRer7 assembly) or *Branchiostoma lanceolatum* (Bl71nemr assembly, see http://amphiencode.github.io and (Marlétaz, et al. 2018)). Reads located in fragments flanked by two restriction sites of the same enzyme, in fragments smaller than 40 bp or within a window of 10 kb around the viewpoint were filtered out. Mapped reads were then converted to reads per first enzyme fragment ends and smoothened using a 30-fragment mean running window algorithm. The smoothened signals were uploaded to the UCSC Genome Browser for visualization.

***Enhancer reporter assays in zebrafish***

To probe the conservation of HCNRs associated with *isl2a* and *isl1* in zebrafish, we cloned the corresponding genomic fragments, using the following primers: *isl1*, 5'-CCGGTTTGGCACTGTATTTA-3' and 5'-TTGGCCCACGTTTGTATAAG-3 to amplify a 908 bp region; *isl2a*, 5'-CTGACAATGCAATAGTGCAG-3' and 5'-TCCATGTATCATCTGAAGGG-3' to amplify an 857 bp region. Similarly, the 2538 bp open chromatin region located between *hey2* and *hddc2* (*hey2-E1)* was amplified with the primers: 5'-CTGGGTTCACAAGCTGACAC-3' and 5'-CATTGGCCCTTGAAACAGAG-3'. The sequences were cloned into the Zebrafish Enhancer Detection (ZED) vector (Bessa, et al. 2009). An injection mix with final concentrations of 40-100 ng/µl plasmid DNA, 50-100 ng/µl Tol2 transposase mRNA, and 0.05% phenol red dissolved in ultrapure distilled $H_20$ (Invitrogen, UK). Concentrations were increased from previous recommendations to gain higher proportions of transgenic embryos without death rates varying from that of uninjected controls. Fertilized AB zebrafish eggs were injected at the 1-2 cell stage using a FemtoJet (Eppendorf, Germany), micromanipulator, and Olympus SZX16 stereomicroscope. As pulled glass capillaries were used to create needles, the pressure and timing of the injection was manipulated to give an approximately 1 nl bolus. F0 larvae were screened for expression of GFP as well as for RFP in the muscle, a marker of integration. Larvae with successful integrations showing a high level of fluorescence were raised to adulthood. At maturity, the F0 mosaic fish were crossed to AB fish, and the F1 offspring screened for expression of GFP and RFP during the first four days of development. Images of reporter fluorescence in zebrafish embryos were collected using cell Sens Entry 1.6 from Olympus Corporation (Japan). At least three founders were obtained for each tested sequence. All protocols used for zebrafish were approved by the Institutional Animal Care and Use Ethic Committee (PRBB–IACUEC) and implemented according to national and European regulations. All experiments were carried out in accordance with the principles of the 3Rs (replacement, reduction and refinement).

**Supplementary Tables**

**Supplementary Table S1 - Ancient GRB pairs present in the last common ancestor of chordates.** Legend: PairID, internal identifier of the GRB pair, including the gene symbols of the linked GRB pairs; N_linked, number of retained GRB pairs in the human genome; N_TD_unliked, number of unlinked *trans-dev* ohnologs in the human genome; N_BS_unlinked, number of unlinked bystander ohnologs in the human genome; TD/BS_linked/unlinked, gene symbols for each gene category; N_NVsp, number of non-vertebrate species for which conservation was observed (maximum number is 8 for source [i, (Irimia, et al. 2012)], 6 for source [ii, (McEwen, et al. 2006)] and 5 for source [iii, this study]); Ancestry_key, indicates the source ([i] GRS, [ii] McE, [iii] NEW) and conservation code; TD_FamilyID, OrthoFinder cluster and ohnolog identifier, Multi_TD/Multi_BS, whether or not multiple copies of the *trans-dev*/bystander gene have been maintained in the human genome; Diff_BS_retention, whether or not the GRB pair is part of an ancestral multi-bystander GRB with differential bystander loss; Extra_Gga/Loc/Cmi, additional GRB pair copies in chicken (Gga), spotter gar (Loc) or elephant shark (Cmi); Single/Multi BS, whether or not the *trans-dev* gene is associated with one or more bystanders.

**Supplementary Table S2 - Ancient coregulated gene pairs in the human genome.** Legend as in Table S1.

**Supplementary Table S3 - Intergenic lengths of *trans-dev* genes compared in Figure 1C.** Legend: TD_ID, GRB pair identifier or arbitrary identifier for "All td" set; Type, LINKED (*trans-dev* gene associated with an ancient bystander in the human genome), LINKED_T-N2 (same as LINKED, but considering the distance of the *trans-dev* [T] and the neighbor gene [N2] after the bystander [B]), LOST_TD (unlinked *trans-dev* gene), TD_ALL (set of all *trans-dev* genes associated with at least one non-*trans-dev* gene); Gene_N1/T/B/N2, Ensembl gene ID for the neighbor next to the *trans-dev* gene (N1), the *trans-dev* gene, and either the bystander gene (for LINKED, LOST_TD or TD_ALL) or the neighbor next to the bystander gene (for LINKED_T-N2); Pos_N1/T/B/N2, linear position of the corresponding gene in the chromosome (from most upstream to most downstream); LENGTH_N1-T/T-B/T-N2, length between N1 and *trans-dev* gene (all cases), between *trans-dev* and bystander genes (for LINKED, LOST_TD or TD_ALL) or between *trans-dev* gene and N2 (for LINKED_T-N2).

**Supplementary Table S4 - Duplicated HCNRs present in a bystander of an ancient GRB and an intergenic region of an unlinked ohnolog.** Legend: Pair ID, GRB pair identifier, as per Table S1; Original_coord (hg17/mm10), coordinate from the element as reported by (McEwen, et al. 2006), in human hg17, or of the ATAC-seq peak in the mouse mm10 genome assembly (in red); BS length, length of the bystander gene body; McEwen ID, identifier used by (McEwen, et al. 2006).

**Supplementary Table S5 – Number of conserved microsyntenic pairs identified for each pair of species.** Using pairwise orthology relationships or clusters from OrthoFinder.

**Supplementary Table S6 - List of human ohnolog relationships compiled for this study.**

**Supplementary Table S7 - List of ATAC-seq experiments used in this study.**

**Supplementary Table S8 - List of primers used in the 4C-seq experiments of this study.**
